# The invasive brown seaweed *Rugulopteryx okamurae* (Dictyotales, Ochrophyta) continues to expand: first record in Italy

**DOI:** 10.1101/2023.09.23.559131

**Authors:** Giancarlo Bellissimo, María Altamirano, Antonio Román Muñoz, Julio De la Rosa, Tin Hang Hung, Gabriele Rizzuto, Salvatrice Vizzini, Agostino Tomasello

**Affiliations:** Regional Agency for Environmental Protection of Sicily (ARPA Sicilia), Complesso Roosevelt, Lungomare Cristoforo Colombosnc, 90149 Palermo, Italy; Universidad de Málaga, Departamento de Botánica y Fisiología Vegetal, Facultad de Ciencias, Campus de Teatinos s/n, 29080 Málaga, Spain; Universidad de Málaga, Departamento de Biología Animal, Facultad de Ciencias, Campus de Teatinos, 29010 Málaga, Spain; Universidad de Granada, Departamento Botánica, Facultad de Ciencias, Campus Fuente Nueva s/n, 18071 Granada, España; Department of Biology, University of Oxford, South Parks Road, Oxford OX1 3RB, UK; National Inter-University Consortium for Marine Sciences, CoNISMa, Piazzale Flaminio 9, 00196 Roma, Italy; Department of Earth and Marine Sciences, University of Palermo, Viale delle Scienze, 90128 Palermo, Italy

**Keywords:** distribution modelling, favorability, fishing, macroalgae, marine bioinvasion, sea current, Sicily

## Abstract

The brown seaweed *Rugulopteryx okamurae* (Dictyotales, Ochrophyta), native to the Pacific Ocean and widely distributed in Asia, has been recently recognized as an emblematic case of biological invasion by marine macroalgae in European waters. Since 2015 and from the Strait of Gibraltar, *R. okamurae* has rapidly spread towards Atlantic and Mediterranean coastal areas exhibiting an invasive behaviour with significant ecological and economic impacts. Here, we report by morphology and genetics the first observation of this species in Italy along the north-western coast of Sicily (Gulf of Palermo), as drifted material and an established population on *Posidonia oceanica*, representing its new eastern distribution limit in the Mediterranean Sea, previously established in Marseilles (France). Furthermore, we have performed with the current introduced distribution of the species a favorability distribution model for the Mediterranean, which shows most of the western Mediterranean, including the Balearic archipelago, Corsica and Sardinia, central Mediterranean, including Sicily, and the northern coast of Africa together with eastern Mediterranean basin, as highly favorable for *R. okamurae*. Arrival of the species into this new area is suggested by means of sea currents and maritime traffic, including fishing activities, hypothesis supported by some of the ranked variables that entered the favorability model, i.e, current velocity, and proximity of fishing ports. These results are a warning that the species can cover large sea distances favored by sea currents, thus also threatening the ecosystems and marine resources of the central and eastern Mediterranean, highly favorable regions for the species. We suggest coordinated actions at the European level regarding prevention, among which those that have the complicity of the fishing sector should be considered, both because it is a highly affected sector and because it potentially has a very important role in the dispersion of the species.

## Introduction

The introduction of alien species into new habitats, mediated by anthropic activities, is one of the greatest threats to the natural capital on which human well-being depends, since some of them can become invasive, causing several-fold impacts on natural and socio-economic systems (Simberloff et al. 2013; Anton et al. 2019; Díaz et al. 2019; Korpinen et al. 2019). Biodiversity is threatened by these species, which pose a continuous challenge for governments trying to minimize introduction events and the impacts they entail. The challenge is even greater in the case of the marine environment due to its spatial and ecological heterogeneity, the absence of barriers, as well as the difficulty of control and study activities, all of which implies an increased difficulty of finding and controlling invasive alien species.

The Mediterranean Sea is a hot spot of introduction of exotic species and invasion events, mainly due to the intense maritime traffic that it entails, as well as its communication with two large bodies of water of a different biogeographical nature (Real et al. 2021). Of the 1001 species introduced on the European coasts, 578 are in the Mediterranean Sea (Zenetos et al. 2022a, b). Furthermore, an upward trend has recently been detected in the rate of appearance of new exotic species, which for the period 2012-2017 has been estimated as an average of 21 species per year (Zenetos et al. 2022a, b). Moreover, 3 out of 4 exotic species that are introduced manage to establish themselves, and 3 out of 4 exotic species are also identified for the first time in Mediterranean waters, mainly originating from the Indo-Western Pacific region (Tsiamis et al. 2018). The Mediterranean Sea can therefore be considered a thermometer for introductions throughout Europe. The introduction vectors seem to be well identified for the marine environment, being ballast water, biofouling and aquaculture activities the most important (Tsiamis et al. 2018). However, the dispersal vectors once the species has been introduced are not as well known or understood, despite the great importance they have in the successful establishment of new species outside their native distribution area, as well as in slowing down their advance and minimizing impacts in the context of control strategies. Thus, understanding the role of ocean currents, recreational aquatic and fishing activities in the dispersal area of species, once they have been introduced may offer important clues on how to curb the expansion of invasive alien species.

Invasive species attract the attention of citizens and governments, because of the sometimes highly visible impacts they produce on natural ecosystems (Katsanevakis et al. 2014; Diagne et al. 2021). From the list of the 10 most invasive species in terms of their negative effects on biodiversity in the Mediterranean Sea, mainly due to competition for resources, the first six are macroalgae species, in the order (Tsirintanis et al. 2022): *Caulerpa cylindracea* Sonder, *Womersleyella setacea* (Hollenberg) RE Norris, *Lophocladia lallemandii* (Montagne) F Schmitz, *Rugulopteryx okamurae* (EY Dawson) IK Wwang, WJ Lee & HS Kim, *Acrothamnion pressii* (Sonder) EM Wollaston and *Caulerpa taxifolia* (M Vahl) C Agardh. Seaweeds are important invasive species as they act as ecosystem engineers with a high capacity to homogenize native communities, with important consequences for trophic chains. Of the six species mentioned above, the one most recently discovered - *Rugulopteryx okamurae* fourth in terms of harmfulness - has also been the first and only species of macroalgae to be included in the list of invasive alien species of Union concern (UE Nº 1143/2014) not only for its detrimental effects on biodiversity, but also for the socio-economic implications of its fast spread, with estimated financial losses of several millions of euros (CIRCABC 2020; MITECO, 2022).

*Rugulopteryx okamurae* (Dictyotales, Ochrophyta)is a subtropical brown seaweed species naturally distributed between the subtropical and temperate western Pacific Ocean (Lee and Kang 1986; Silva et al. 1987; Yoshida 1998). It exhibits a diplobiontic isomorphic life cycle, common to other morphologically similar species of the same family, as *Dictyota* or *Taonia* spp., which makes its introduction a bookcase of cryptic invasion. Furthermore, in its native area, the species exhibits up to three different morphotypes along the year (Sun et al. 2006), which seems to occur as well in the newly introduced alien populations (Salido and Altamirano 2020). As in the native populations, the species seems to fail the sexual reproduction, maintaining and establishing new populations through vegetative (propagules) and asexual (monospores) reproduction (Hwang et al. 2009; Altamirano et al. 2016, 2017), throughout the year (Salido and Altamirano 2020). The first record outside its native area was in Thau Lagoon in France in 2002 (Verlaque et al. 2009), where the species is present but not invasive (Ruitton et al. 2021). The first record as an invasive species was on the Spanish coast of the Strait of Gibraltar in 2016 (Altamirano et al. 2016, 2017), on the African coast (Ceuta, Spain) and then on the European coast. Since then, the species has been spreading both towards the Mediterranean Sea and the Atlantic Ocean, being already present other than in Spain (Altamirano et al. 2016; García Gómez et al. 2020), in Morocco (El Aamri et al. 2018), Portugal (Liulea et al. 2023) and France (Ruitton et al. 2021), having even reached different Macaronesian islands such as the Azores (Faria et al. 2021), Madeira (Bernal-Ibáñez et al. 2022) and the Canary Islands (REDEXOS 2022). Its limit in the Mediterranean is currently found on the coast of Marseilles, in France (Ruitton et al. 2021). The species has a high capacity for vegetative proliferation as well as for dispersal, which gives it a high capacity to produce large amounts of biomass that can be moved by currents (García-Lafuente et al. 2023; Mateo-Ramírez et al. 2023) or helped by human vectors such as fishing or maritime transport (MITECO 2022), and which can end up forming important wracks on the coasts. The seaweed’s ability to fast proliferation is causing serious concern, because it greatly impacts both native natural communities (seagrasses and marine forests), reducing their extension and diversity (García Gómez et al. 2020, 2021), and human activities such as fishing and tourism, with losses of millions of euros across various sectors (CIRCABC 2020; MITECO 2022). *R. okamurae* sparked major concerns first only in Spain and Morocco, but since 2022 it has been recognized as threat to marine ecosystems by the European Union, with its inclusion in the European List of invasive species (EU 2022/2013). The Mediterranean Sea has been identified as a highly favourable habitat for this species (Muñoz-Gallego et al. 2019), which points to potential increasing impacts if its spread on European coasts is not stopped. It is therefore essential to set up an early warning system for this species, by identifying the dispersal vectors and the entry points in new areas and elaborate coordinated protocols for its control.

In this context, the objective of this work is to report the first record of *R. okamurae* in Italy, on the island of Sicily, being the easternmost of the species in the Mediterranean Sea to our knowledge and providing the current favorability for the species in the Mediterranean. This work provides additional evidence of the fast dispersal of this species in the eastern Mediterranean Sea, an area highly favorable to its establishment, especially under the current climate scenario.

## Materials and Methods

### Study site

Massive beached monospecific seaweed biomass was observed for first time from the end of June 2023 on the shore of Aspra (38°06’17” N 13°29’52” E), a small fishing village within the eastern side of the Gulf of Palermo along the north-western coast of Sicily (Figure 1), on rocks and concrete blocks scattered along the shore (Figure 2). The coastal zone facing the stranding area has sandy bottoms also featured by dead vermetid reefs and rocky blocks at shallow depths, while artificial submerged and subaerial breakwaters run parallel to the shoreline at about 80 m of metres from the shoreline. At the beginning of July, more drifted material was found in a locality adjacent to the main port area of Palermo (38°07’14” N - 13°22’18” E) and at the end of August on the beach of the Port of Bandita (38°05’54” N - 13°25’01” E), where detritus was observed on rocks and concrete blocks scattered along the shore, and at the end of August on the beach of the Port of Bandita (38°05’54” N - 13°25’01” E) (Figure 2). An established population of *R. okamurae* was found in July 2023 in Aspra on *Posidonia oceanica* (L.) Delile plants at 4 m depth, with high density, for which *in situ* observations and relative abundance of *R. okamurae* were performed in an area of about 1 ha. A further monitoring for the species was performed in July and August 2023 along the coast of Sicily (Figure 2) and Lampedusa Island to study the distribution of the species (Figure 1 and Supplementary Table S1).

**Figure 1.**
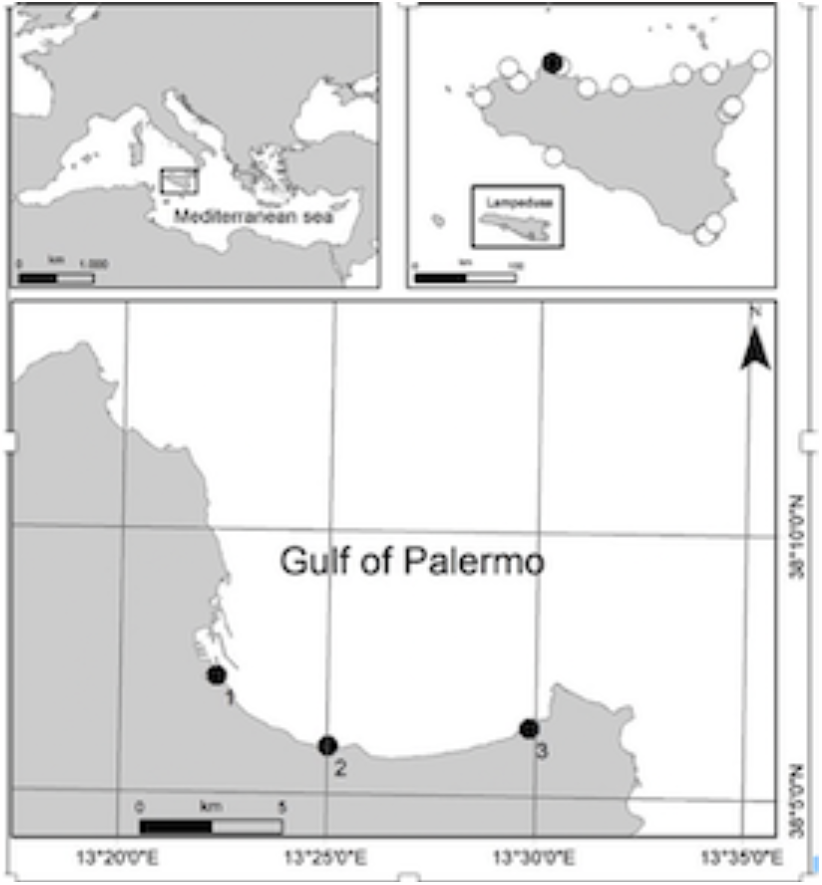
Distribution of *R. okamurae* found along coast of Sicily and Lampedusa Island in summer 2023. Black dots indicate the presence of the species (1) Port of Palermo, (2) Port of Bandita, (3) Aspra beach, white dots indicate visited places where the species was not found.

**Figure 2.**
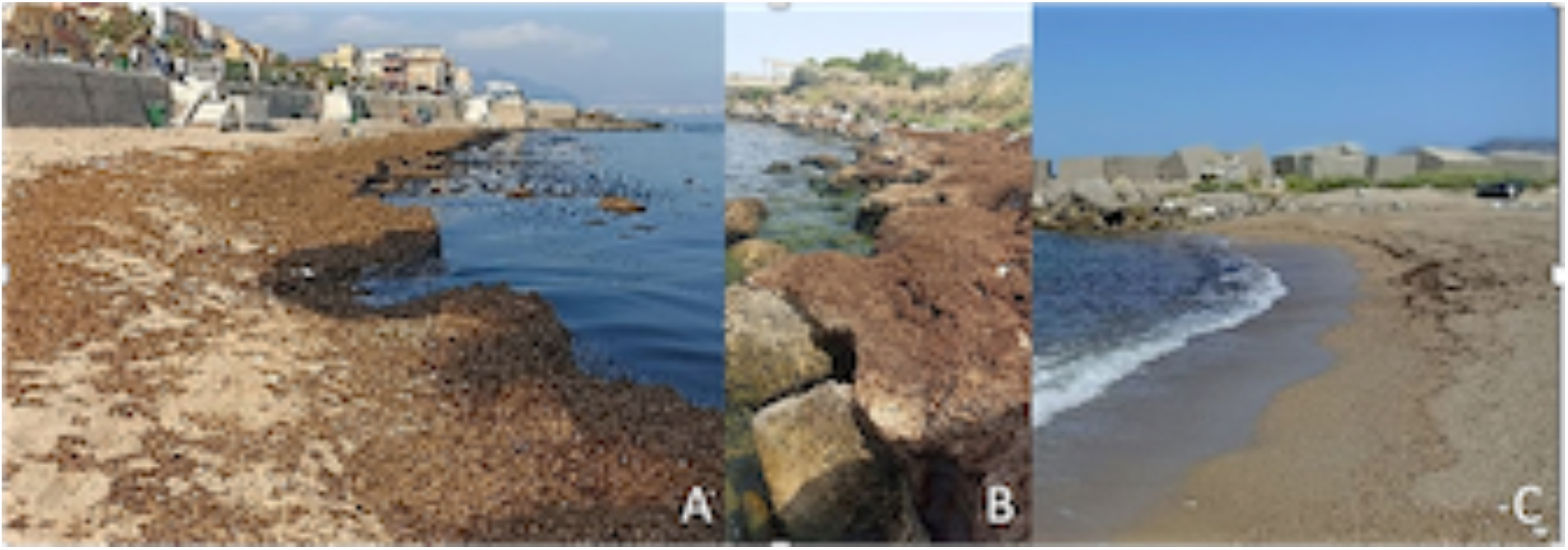
Drifted biomass of *R. okamurae* on the beach of Aspra (A), on artificial rocky blocks at Port of Palermo (B) and along the shoreline at Port of Bandita (C).

### Collection and identification

Specimens were collected in all discovery sites, put into ziplock bags, and stored under dark and cold conditions until morphological identification was performed in the laboratory. Samples from drifted material were collected from the outermost front of the mass and those from established underwater population were collected at depths ranging from 0.2 to 4 m. In the laboratory, thalli were observed using both a 3D Digital Microscope (Hirox RH-2000) and a stereomicroscope (Discovery V20, Zeiss). Morphological identification was performed on material collected at all sites in accordance with reference literature (Womersley 1987; De Clerck et al. 2006; Sun et al. 2006; Hwang et al. 2009). Voucher specimens were deposited in the personal herbarium of GB at the Regional Agency for Environmental Protection in Palermo (ARPA Sicilia – UOC Area Mare) and in University of Málaga herbarium (MGC-Phyc 5515).

Three samples were collected for species-level identification by DNA analysis and put into silica gel. Genomic DNA extraction was performed using a *DNeasy Plant Mini Kit (Qiagen, United Kingdom)*, following the manufacturer’s instructions. DNA yield and quality were assessed by using a NanoDrop One Spectrophotometer (Thermo Fisher Scientific, United Kingdom). Only two samples passed the quality check and were retained for downstream work. Genes encoding chloroplast proteins rbcL and psbA were amplified by PCR using a ProFlex PCR System (Thermo Fisher Scientific). The primers were based on a previous study of Faria et al. (2022), with modifications to primers targeting the rbcL locus (see Supplementary Table S2). The 25-μl reactions contained: 12.5 μl Q5 Hot Start High-Fidelity 2X Master Mix (New England Biolabs, United Kingdm), 1.25 μl forward primer (10 μM), 1.25 μl reverse primer (10 μM), and 10 μl genomic DNA (∼ 50 ng). The thermal cycling profile was: 98°C 30 s, 30 × [98°C 30 s, 60°C 30 s, 72°C 1 min], 72°C 2 min. PCR product size and integrity were verified by electrophoresis on 1% agarose gel. Reactions were purified using a *Monarch* PCR & *DNA Cleanup Kit (New England Biolabs)* and sent to Source BioScience, Nottingham, United Kingdom for Sanger sequencing in both forward and reverse directions. Nucleotide sequences were inspected and trimmed to remove low base-call quality reads on Unipro UGENE v48.0. Finally, DNA sequences were aligned on MAFFT and the consensus sequence was uploaded on BLASTn for multiple pairwise comparisons with reference sequences stored at the NCBI database.

### Distribution modelling

To model the distribution of *R. okamurae* we used all known records in the Mediterranean by September 2023 (Figure 3), which were restricted to the western Mediterranean, until the appearance of the species in Italy. In the modelling procedure, we used the Bio-ORACLE database (version 2.1 and 5 arc minutes resolution), which consists of a set of geophysical, biotic, and climatic data for the oceans of the planet (Assis et al. 2018). We also used two variables related to the distance to fishing and commercial harbours. The distribution data of *R. okamurae* and the environmental layers were projected onto a hexagonal grid, with each hexagon representing Operative Geographical Units (OGU) stretching an area of 7774 km^2^. To control multicollinearity among environmental variables we calculated Spearman correlation coefficients and for each pair of variables with r > 0.8, we only retained the variable with the highest individual predictive power (Chamorro et al. 2021, García-Carrasco et al. 2023). We limited the increase in type I error bound by the number of variables analyzed by addressing a False Discovery Rate control (Benjamini and Yekutieli 2001).

**Figure 3.**
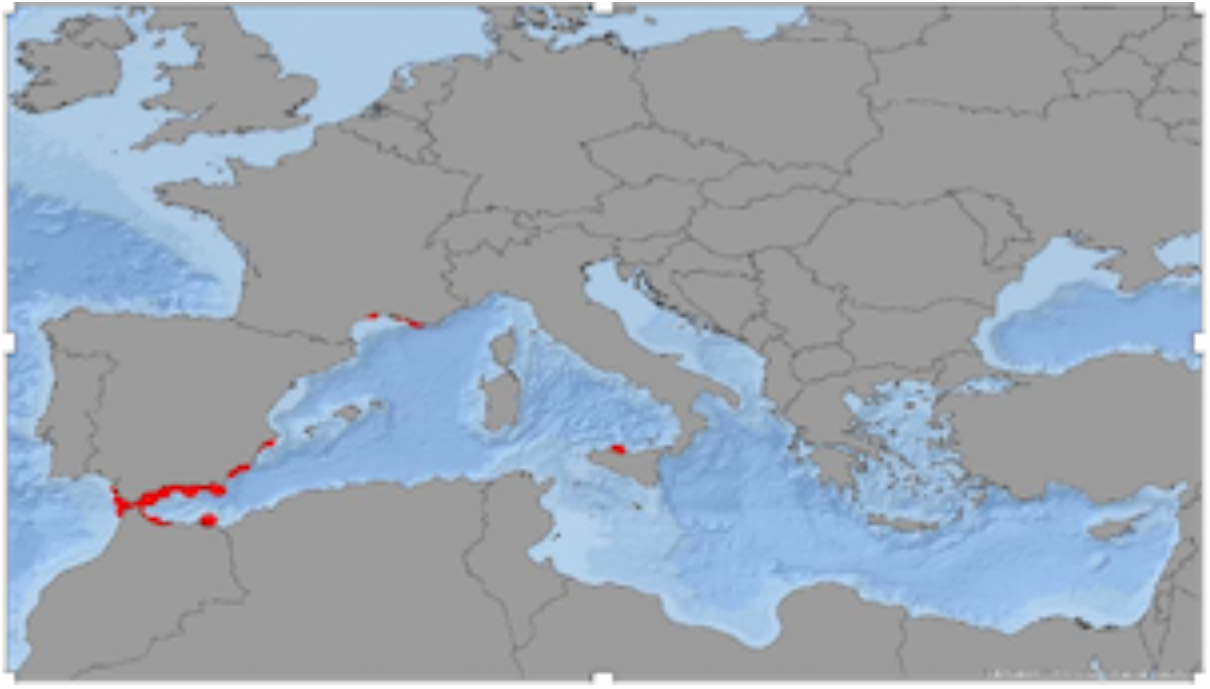
Current Mediterranean distribution of *R. okamurae*.

Considering that the risk of invasion derives from the degree to which the environmental conditions are favorable for the presence of the species we build a risk model for *R. okamurae* invasion in the Mediterranean by performing a multivariable forward-backward stepwise logistic regression of the current presences/absences on the set of variables, in a way that variables were added to an initial null model if their inclusion significantly improved the regression. Based on the probability values resulting from the logistic regression we obtained the favorability values, ranging from 0 to 1, by applying the Favorability Function (Muñoz et al. 2005; Real et al. 2006). These values represent deviations from the expected chance of *R. okamurae* invasion and identify localities with environmental conditions associated with already invaded areas. We used the Wald test to assess the weight of each variable and the estimation of the parameters in the logit equation (Wald 1943). To evaluate the discrimination capacity of the model we used the Area Under the Receiver Operating Characteristic Curve (AUC; Hosmer and Lemeshow 2000). This modelling approach complies with standard modelling protocols (Sillero et al. 2021).

## Results

Morphology and anatomy of collected samples at all sites corresponded to the invasive brown seaweed *R. okamurae*. External vegetative morphology of thalli was flat ribbon-like corrugated and dichotomous branched up to 18 cm height with stoloniferous attaching system, exhibiting a yellow-brown colour without iridescence (Figure 4). Inner structure of the thalli was characterized by a monostromatic cortical, a monostromatic central medulla, and a pluristromatic medulla at the margin of the thalli (Figure 5). Two different morphotypes with this general anatomical pattern were observed in collected samples. Drifted material on the beach exhibited a thick morphotype, with more than 4 medullar cells at the margin of the thalli and wide branches up to 1 cm (Figure 5A). Thalli from established population exhibit a thin morphotype with 2-3 medullar cells at the margin of the thallus and narrow branches (<0,5 cm) (Figure 5B). Vegetative propagules were observed in specimens from all sites, but no asexual monospores or sexual gametangia. Some epiphytic red algae, such as encrusting coralline *Hydrolithon farinosum* (JV Lamouroux) Penrose & YM Chamberlain, articulated coralline *Jania rubens* (Linnaeus) JV Lamouroux and filamentous *Ceramiales* were detected entangled or epiphyte on *R. okamurae*. No evidence of grazing was observed.

**Figure 4.**
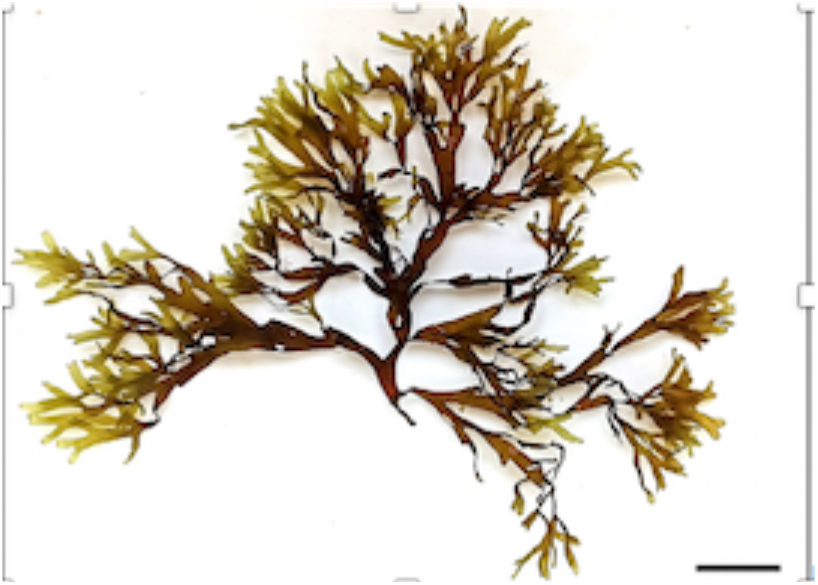
*Rugulopteryx okamurae* specimen sampled in the established underwater population of Aspra. Scale bar = 3 cm.

**Figure 5.**
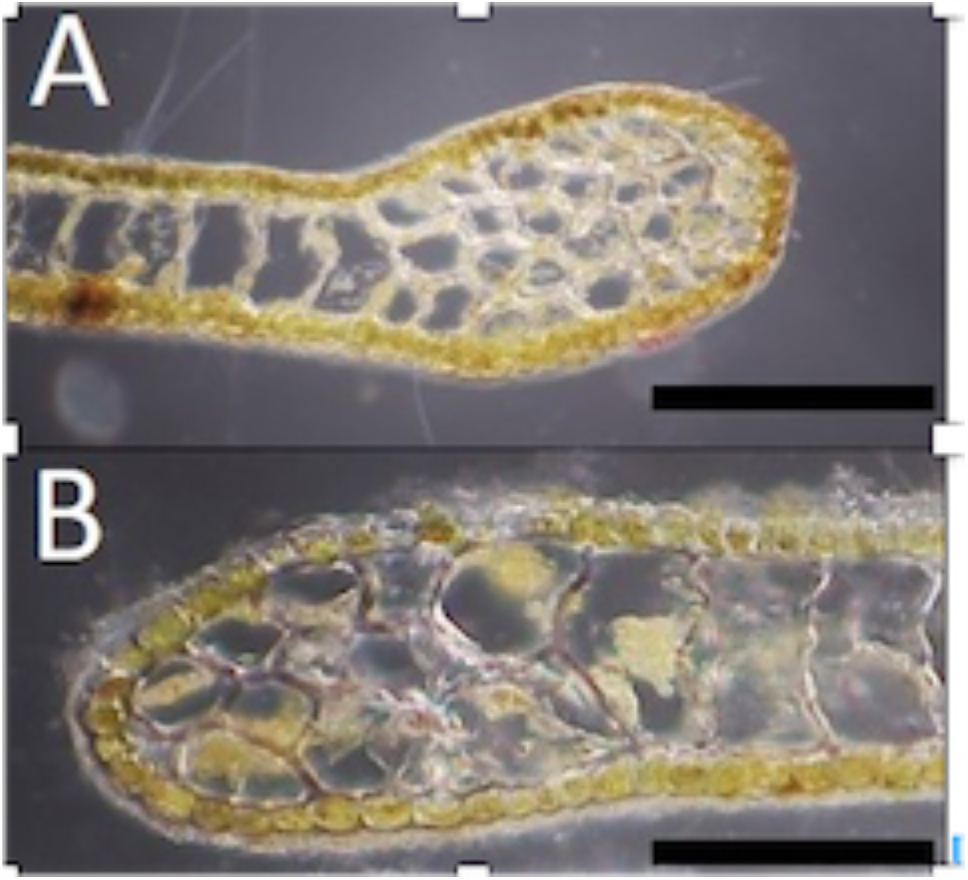
Cross-section of a specimen both beached (A) and from the settled population (B) showing a multi-layered medulla near the margin. Scale bar = 200 μm.

BLAST searches of psbA and rbcL gene sequences returned 99% to 100% identity scores compared to *R. okamurae* specimens collected around the world, including Madeira and Azores (Bernal-Ibáñez et al. 2022; Faria et al. 2022), Cádiz and Alicante (Spain), and the native areas of Korea and Japan (see Supplementary Table S3 and S4). Overall, these results confirmed the morphological identification at the species-level of the samples collected along the sicilian coast as *R. okamurae* thalli. The newly obtained seqeuences were deposited in NCBI GenBank with the accession numbers OR578566–OR578567.

The stablished population of *R. okamurae* was found invading small *P. oceanica* patches extending about 100 m^2^ at a depth of 3-4 m. The invasive seaweed was epiphyte on both rhizome and leaves of *P. oceanica*, competing with native flora (Figure 6A). Attached thalli of *R. okamurae* were also observed on breakwaters (Figure 6B), sharing habitat with other macroalgae species such as *Asparagopsis taxiformis* (Delile) Trevisan, *Laurencia* complex and articulated corallines *Ellisolandia elongata* (J Ellis & Solander) K Hind & GW Saunders and *Jania rubens*, exhibiting a coverage of approximately 60% with a depth range of 2-4 m. Water temperature at the collection moment was 27 ºC.

**Figure 6.**
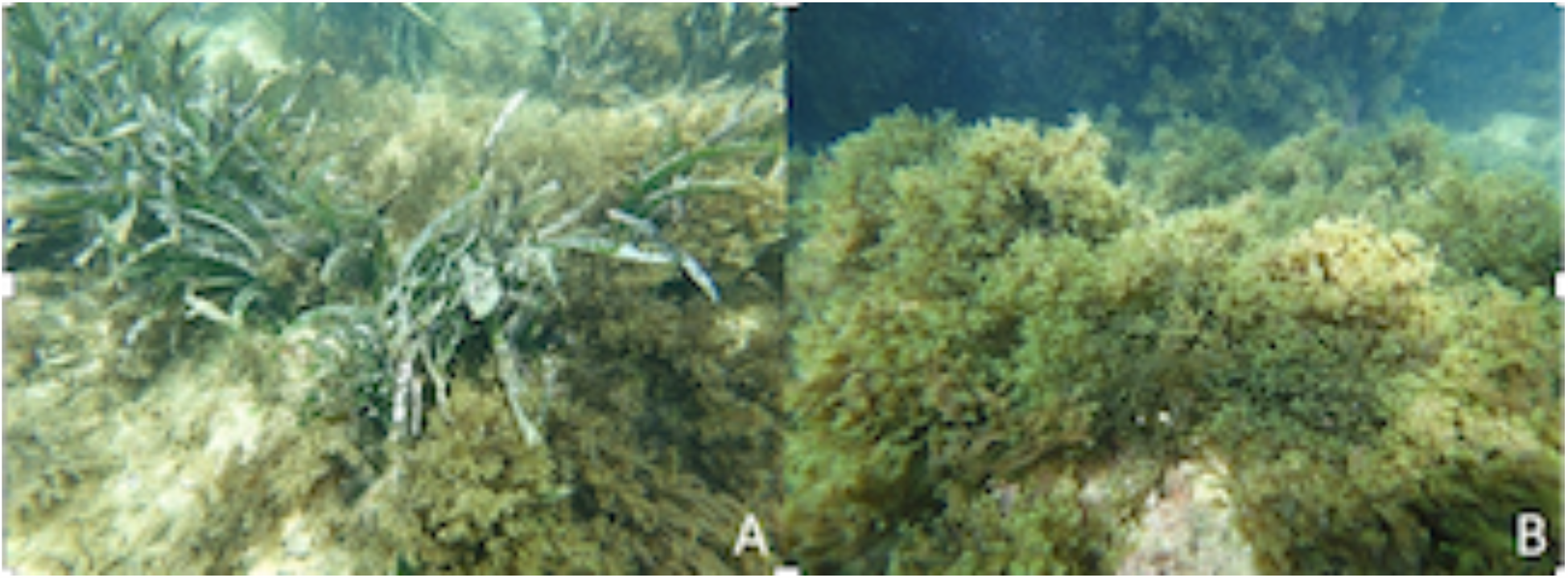
A) Established population of *R. okamurae* on *P. oceanica* and B) with native algal flora on artificial rocky blocks.

We obtained a significant environmental favourability model with high discrimination capacity (AUC > 0.88). This model shows most of the western Mediterranean, including the Balearic archipelago, Corsica and Sardinia, central Mediterranean, including Sicily, and the northern coast of Africa together eastern Mediterranean basin, as high favorable for *R. okamurae* (Figure 7). Variables that entered and explained the model are more related with vectors than with factors determining biological activity of the species and are ranked the following way: mean and minimum currents velocity, cloud cover, phosphate, and distance to fishing harbours (see Supplementary Table S5).

**Figure 7.**
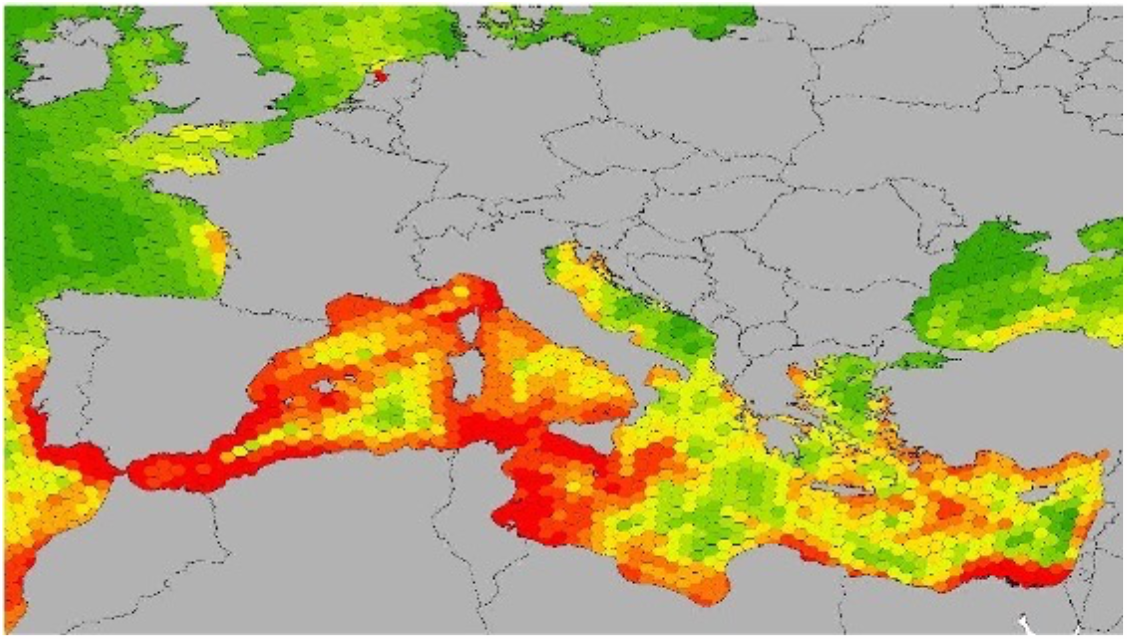
Cartographic representation of the current environmental favourability for *R. okamurae* in the Mediterranean in each operational geographic unit.

## Discussion

The morphological and genetic evidence provided in this work confirms the expansion of the invasive seaweed *R. okamurae* towards the Mediterranean Sea and its introduction in Italy. Chloroplast genome sequences are widely used for plant and algal species discrimination (Costa et al. 2016; Sun et al. 2020; Liu et al. 2022; Song et al. 2023). The genes coding for the photosystem reaction center protein D1 (psbA) and the ribulose-1,5-bisphosphate carboxylase/oxygenase large subunit (rbcL) are molecular markers commonly used for brown seaweed phylogenetic studies (Saunders and Moore 2013). Here, we used primers targeting these chloroplast genomic regions to corfirm morphological observations and obtained accurate identification at the species level (99%-100%) of the samples as the brown seaweed *R. okamurae*. This molecular analysis proved a close relationship with specimens previously collected in the Atlantic Ocean (Faria et al. 2022, Bernal-Ibáñez et al. 2022).

The species has once again shown its ability to cross great distances of open sea to reach islands, in this case the island of Sicily in Italy. Previously, the species managed to reach the Macaronesian islands, first Azores (Faria et al. 2021, 2022), and afterwards Madeira and the Canary Islands (Bernal-Ibáñez et al. 2022; REDEXOS 2022), also covering great distances in open sea. The presence of *R. okamurae* in Sicily may have two explanations that are not exclusive, but rather, on the contrary, synergistic. On the one hand, the species may have arrived in the form of large accumulations of biomass transported by currents, as already observed in the Alborán Sea (García-Lafuente et al. 2023; Mateo-Ramírez et al. 2023) and suggested to explain the presence of the species in Madeira (Bernal-Ibáñez et al. 2022). In other words, marine currents may be facilitating the long-distance dispersal of the species, allowing the arrival of large amounts of biomass that either have been detached from populations established in remote areas, or that have grown in suspension in the water column until it was dumped on the beach. The results of the present work show that this is what may have happened in Sicily too. In fact, wracks were found with thalli exhibiting the thick morphotype in July, while this morphotype is typical of cold months (Sun et al. 2006; Salido and Altamirano 2020), indicating that these thalli may have grown at another time and/or place with lower temperature, probably arriving from some other part of the western Mediterranean, aided by a regime of marine currents that would be interesting to investigate. On the contrary, the thallus of the established population found in the study area presented a fine morphotype, typical of the warmer months (Sun et al. 2006; Salido and Altamirano 2020), coinciding with the time of its finding. The important role of sea currents as natural dispersal vectors for secondary introductions of *R. okamurae* can be inferred as well from favorability model performed from current introduced distribution of the species. Current velocity, expressed as mean and minimum values, are the variables entering the model that more contribute to explain *R. okamurae* distribution. This vector is worrying from the point of view of the management of the species, since it represents a dispersion that is difficult if not almost impossible to control and with an enormous geographical reach inside and outside the Mediterranean.

The other explanation for the arrival of *R. okamurae* in Sicily, potentially synergistic and not exclusive with the previous one, is based on the hypothesis of the arrival of thalli mediated by vectors linked to anthropogenic activities, potentially maritime transport or fishing or recreational activities. This argument was already used to explain the introduction of *R. okamurae* in the Azores islands (Faria et al. 2021), suggesting ballast water and/or fouling as potential vectors, since the species can survive up to three weeks in the dark (Rosas-Guerrero et al. 2018) and it is capable of adhering to different substrates (García-Gómez et al. 2018). The fact that the first records of *R. okamurae* in Sicily have been found very close to an important port strongly support this explanation. The species could have been introduced through ballast water from other points in the Mediterranean that share shipping routes with the port of Palermo or may be the destination of fishing activities carried out in other invaded areas of the Mediterranean, like Marseilles, where *R. okamurae* is well established (Ruitton et al. 2021). Also in this case, the results of the favorability model support this second hypothesis, since proximity to fishing ports, although it was the last variable to enter the model, contributes to explaining the current distribution of the species. Transport-stowaways (attributed to maritime traffic) have been identified as responsible of around half of Italian marine introductions of alien species, and Sicily is a hotspot due to its geographic location as a crossroad between Indo-Pacific and Atlantic waters (Servello et al. 2019). Furthermore, the port of Palermo is one of the major Mediterranean cruise ports serving the passenger shipping traffic mainly between Italy, France, and Greece, linking the island to major Mediterranean cities (Rome, Genoa, Naples, Marseilles, Barcelona, Valencia, Athens and Tunis), besides the tourist marina serving yachts and catamarans.

In either one of the possible introduction scenarios, it would be interesting to know whether *R. okamurae* has reached the coasts of Corsica and Sardinia, as well as the coast of Tunisia, indicating a broad and fast expansion in the Mediterranean, as already suggested by environmental favorability model performed in this study. Regardless the presence of *R. okamurae* on the northern coast of Sicily should be considered as a warning that the species is at the gateway to the central Mediterranean and is advancing towards its eastern part, which is highly favorable for the establishment of this species.

At all sites where *R. okamurae* has been introduced outside its native range since 2016, the species has produced unprecedented environmental and socioeconomic impacts (Altamirano et al. 2016, 2017; El Aamri et al. 2018; García-Gómez et al. 2018; Faria et al. 2021, 2022; Ruitton et al. 2021; Bernal-Ibáñez et al. 2022; REDEXOS 2022; Liulea et al. 2023). The results of the present work do not allow us to assess the entity of the impact of *R. okamurae* on the study region, as it seems that the seaweed is at the beginning of its invasion and the area was already highly degraded by anthropogenic pollution. However, it cannot be ruled out that the introduction of *R. okamurae* in Sicily could have similar impacts to those already observed in other areas, for example on the community associated with *P. oceanica* or on communities of macroalgae (García-Gómez et al. 2018, Faria et al. 2022) and coralligenous (Navarro-Barranco et al. 2021). The presence of the drifted biomass of *R. okamurae* near the port of Palermo should alert the local councils that they can cause complaints from beach users if they are not removed and warn them to reserve budgetary items for this. Furthermore, we want to warn that if the first manifestations of the presence of *R. okamurae* occur as thalli intermingled in banquettes of dead *P. oceanica* leaves, or as wracks that are very similar in appearance to *P. oceanica* banquettes, it can make its detection and therefore early warning difficult. Hence the need to inform and educate citizens and managers about this new invasion. In any case, the consequences of the introduction of *R. okamurae* in Sicily should be studied in detail as a sample of what the species could cause if it spreads towards the eastern Mediterranean.

In the central and eastern Mediterranean, the most important invasive macroalgal species to date is *C. cylindracea*, a species that was present in the Strait of Gibraltar when and where *R. okamurae* began its invasion, together with *Asparagopsis armata* Harvey and *A. taxiformis* (Altamirano et al. 2019). It is possible that extant invasive species may support *R. okamurae* expansion through a process of invasional meltdown (Simberloff and Von Holle 1999), but later may gradually reduce their presence, although they do not completely disappear. The same could happen in those areas of the Mediterranean that are currently heavily invaded by *C. cylindracea*, fulfilling the saying that another will come that the previous better will become.

Management options for *R. okamurae* are challenging at the local, national, and European level, but it should not be a reason for inaction. In this regard, the Spanish Government published in 2022 the national control strategy for *R. okamurae* (MITECO 2022). This document highlights that the prevention measures against the seaweed’s introduction and spread are the key for its control, pointing out the need to identify its dispersal vectors. In this context results of the favorability model performed in this study can be of much help. The favorability map presented indicates favorable areas for *R. okamurae* that have already been invaded, but also areas in which the species has not yet been recorded. Furthermore, the model has identified a variable that points towards a vector of anthropogenic origin, the proximity of fishing ports, and therefore susceptible to management to avoid or minimize the dispersal of the species. And these two results extracted from the model represent an opportunity to work on preventing the introduction of the species, especially in areas of special interest for the conservation of marine habitats. In this context of opportunity to invest efficiently in preventing the introduction of *R. okamurae* into new areas, having the complicity and support of the fishing sector is crucial, and hence the need to inform them about the role they play in the management of this invasive species, not only as an affected sector but also as part of the solution. For this reason, the Spanish control strategy for *R. okamurae* points out the need to develop disinfection protocols for fishing gear, for example, as well as to work together with port authorities and the fishing sector (MITECO 2022). At the European level, there is an urgent need for coordination actions that slow down the dispersal of the species, and that minimize and mitigate its environmental impact on the affected sectors, such as fishing.

## Acknowledgements

We sincerely thank Prof. Amelia Gómez Garreta, from Universitat de Barcelona (Spain) for putting the Italian and Spanish research groups in contact, making this publication possible.

## Supplementary tables

**Supplementary Table S1.** Distribution records of *R. okamurae* in Italy.

| Location |  |  |  |
| --- | --- | --- | --- |
| Mondello | 38°11'58" N | 13°19'45" E | Not observed |
| Port area of Palermo (Porto Borbonico) | 38°07'14" N | 13°22'18" E | Drifted material on artificial rocky blocks |
| Port of Bandita | 38°05'54" N | 13°25'01" E | Drifted material on the shoreline |
| Aspra | 38°06'17" N | 13°29'52" E | On *Posidonia oceanica* |
| Sant'Elia | 38°05'43" N | 13°32'22" E | Not observed |
| Trabia | 38°00'23" N | 13°38'25" E | Not observed |
| Cefalù | 38°02'06" N | 14°00'52" E | Not observed |
| Settefrati | 38°01'49" N | 13°57'52" E | Not observed |
| Marinello | 38°08'26" N | 15°03'17" E | Not observed |
| Capo d'Orlando | 38°08'14" N | 14°43'04" E | Not observed |
| Ganzirri | 38°15'25" N | 15°36'47" E | Not observed |
| Sant'Alessio Siculo | 38°15'25" N | 15°36'47" E | Not observed |
| Taormina | 37°55'01" N | 15°20'43" E | Not observed |
| Giardini Naxos | 37°50'02" N | 15°16'19" E | Not observed |
| Fiumefreddo | 37°47'06" N | 15°14'07" E | Not observed |
| Vendicari | 36°47'59" N | 15°05'39" E | Not observed |
| Porto Ulisse | 36°41'47" N | 14°59'21" E | Not observed |
| Santa Maria del Focallo | 36°41'39" N | 14°57'54" E | Not observed |
| Eraclea | 37°23'19" N | 13°16'28" E | Not observed |
| San Teodoro | 37°54'17" N | 12°27'16" E | Not observed |
| San Vito Lo Capo | 38°10'33" N | 12°44'41" E | Not observed |
| Fossa dello Stinco | 38°02'59" N | 12°51'53" E | Not observed |
| Cala degli Alberelli | 38°03'58" N | 12°49'37" E | Not observed |
| Guidaloca | 38°03'22" N | 12°50'28" E | Not observed |
| Lampedusa island (Rabbit island) | 35°30'48" N | 12°33'26" E | Not observed |
| Lampedusa island (Guitgia) | 35°29'56" N | 12°35'56" E | Not observed |
| Lampedusa island (Mare morto) | 35°30'57" N | 12°37'37" E | Not observed |

**Supplementary Table S2.**
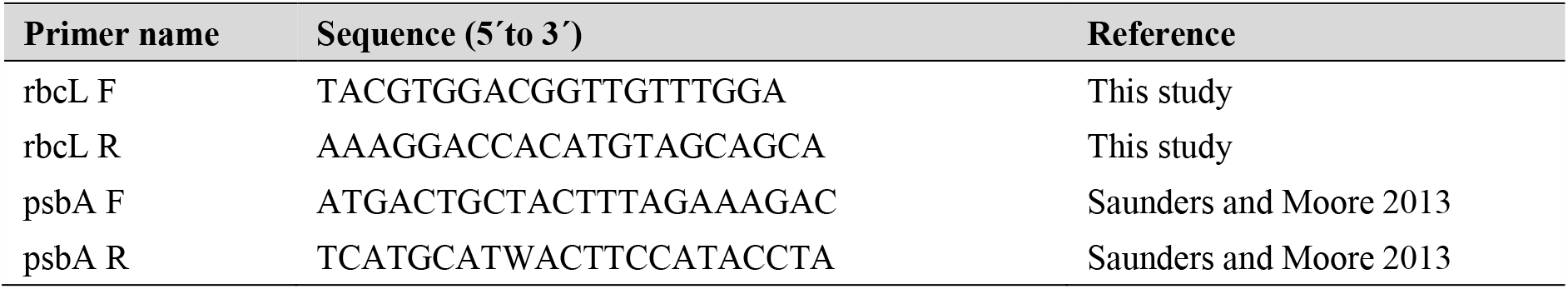
List of primers targeting the genetic markers used in this study.

**Supplementary Table S3.**
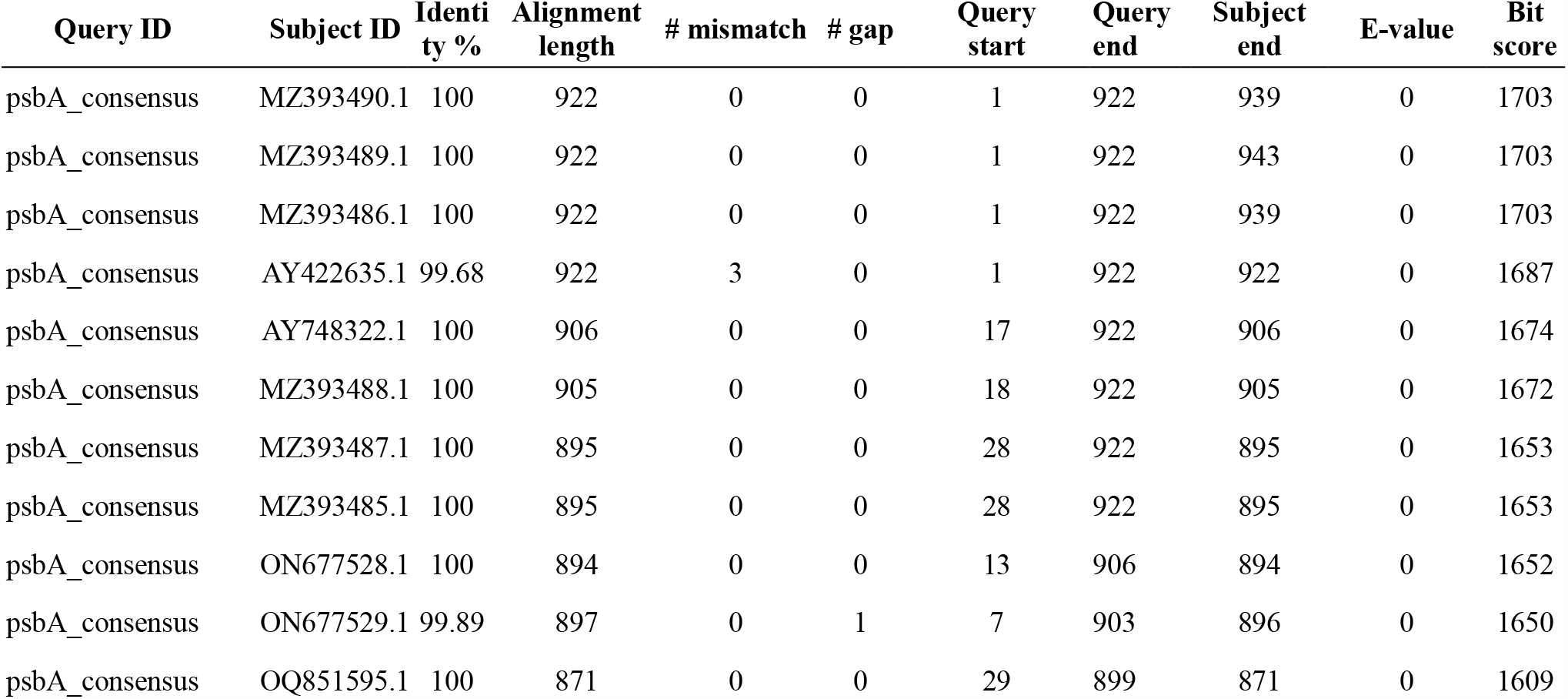
BLAST hit score table for psbA gene sequence searches against the NCBI.

**Supplementary Table S4.**
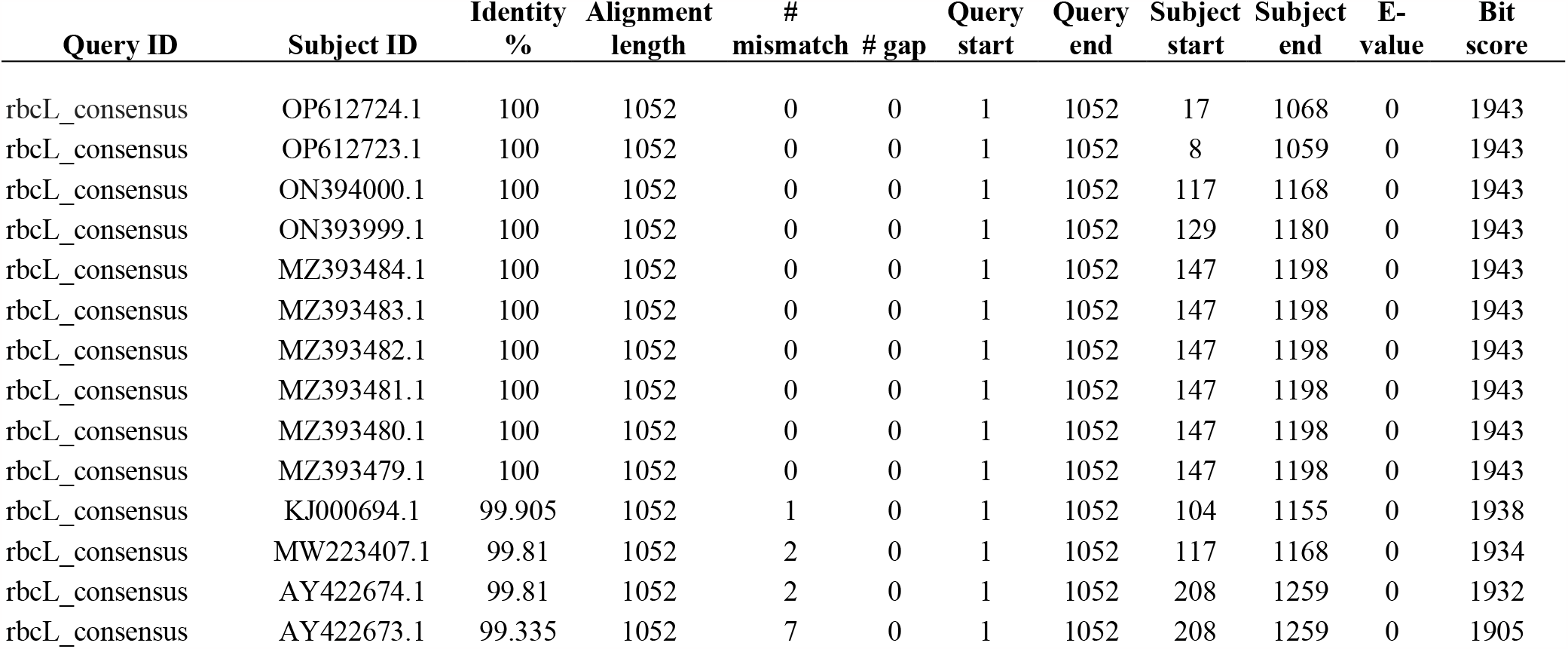
BLAST hit score table for rbcL gene sequence searches against the NCBI database.

| Query ID | Subject ID | Identity % | Alignment length | # mismatch | # gap | Query start | Query end | Subject start | Subject end | E-value | Bit score |
| --- | --- | --- | --- | --- | --- | --- | --- | --- | --- | --- | --- |
| rbcL_consensus | OP612724.1 | 100 | 1052 | 0 | 0 | 1 | 1052 | 17 | 1068 | 0 | 1943 |
| rbcL_consensus | OP612723.1 | 100 | 1052 | 0 | 0 | 1 | 1052 | 8 | 1059 | 0 | 1943 |
| rbcL_consensus | ON394000.1 | 100 | 1052 | 0 | 0 | 1 | 1052 | 117 | 1168 | 0 | 1943 |
| rbcL_consensus | ON393999.1 | 100 | 1052 | 0 | 0 | 1 | 1052 | 129 | 1180 | 0 | 1943 |
| rbcL_consensus | MZ393484.1 | 100 | 1052 | 0 | 0 | 1 | 1052 | 147 | 1198 | 0 | 1943 |
| rbcL_consensus | MZ393483.1 | 100 | 1052 | 0 | 0 | 1 | 1052 | 147 | 1198 | 0 | 1943 |
| rbcL_consensus | MZ393482.1 | 100 | 1052 | 0 | 0 | 1 | 1052 | 147 | 1198 | 0 | 1943 |
| rbcL_consensus | MZ393481.1 | 100 | 1052 | 0 | 0 | 1 | 1052 | 147 | 1198 | 0 | 1943 |
| rbcL_consensus | MZ393480.1 | 100 | 1052 | 0 | 0 | 1 | 1052 | 147 | 1198 | 0 | 1943 |
| rbcL_consensus | MZ393479.1 | 100 | 1052 | 0 | 0 | 1 | 1052 | 147 | 1198 | 0 | 1943 |
| rbcL_consensus | KJ000694.1 | 99.905 | 1052 | 1 | 0 | 1 | 1052 | 104 | 1155 | 0 | 1938 |
| rbcL_consensus | MW223407.1 | 99.81 | 1052 | 2 | 0 | 1 | 1052 | 117 | 1168 | 0 | 1934 |
| rbcL_consensus | AY422674.1 | 99.81 | 1052 | 2 | 0 | 1 | 1052 | 208 | 1259 | 0 | 1932 |
| rbcL_consensus | AY422673.1 | 99.335 | 1052 | 7 | 0 | 1 | 1052 | 208 | 1259 | 0 | 1905 |

**Supplementary Table S5.** Variables that entered the model via a forward–backward stepwise selection process, ranked by their order of entrance. βs are the coefficients in the logit function, S.E. is the standard error of these coefficients, Wald is the Wald’s statistics value (representing the relative importance of the variable in the model) and p is the significance of the coefficients according to the Wald test.

| Variable | $\beta$ | S.E. | *Wald* | *p* |
| --- | --- | --- | --- | --- |
| Maximum cloudiness | -0.962 | 0.299 | 10.338 | 0.001 |
| Mean current velocity | 26.334 | 3.410 | 59.631 | <0.001 |
| Minimum velocity current | -10.617 | 2.380 | 19.892 | <0.001 |
| Phosphate range | -7.012 | 2.649 | 7.006 | 0.008 |
| Proximity to fishing port | 0.027 | 0.015 | 3.385 | 0.066 |
| Constant | -5.220 | 0.697 | 56.056 | <0.001 |

